# The PPLD has advantages over conventional regression methods in application to moderately sized genome-wide association studies

**DOI:** 10.1101/2021.05.03.442456

**Authors:** Veronica J. Vieland, Sang-Cheol Seok

## Abstract

In earlier work, we have developed and evaluated an alternative approach to the analysis of GWAS data, based on a statistic called the PPLD. More recently, motivated by a GWAS for genetic modifiers of the X-linked Mendelian disorder Duchenne Muscular Dystrophy (DMD), we adapted the PPLD for application to time-to-event (TE) phenotypes. Because DMD itself is relatively rare, this is a setting in which the very large sample sizes generally assembled for GWAS are simply not attainable. For this reason, statistical methods specially adapted for use in small data sets are required. Here we explore the behavior of the TE-PPLD via simulations, comparing the TE-PPLD with Cox Proportional Hazards analysis in the context of small to moderate sample sizes. Our results will help to inform our approach to the DMD study going forward, and they illustrate several respects in which the TE-PPLD, and by extension the original PPLD, offer advantages over regression-based approaches to GWAS in this context.

## 1. Introduction

In previous work we have developed and evaluated a statistic called the Posterior Probability of Linkage Disequilibrium (PPLD) as a measure of evidence for or against trait-SNP association (1–3), and we have extended the PPLD to accommodate time-to-event (TE) data, yielding the TE-PPLD (4). In this paper we compare and contrast the TE-PPLD with the more familiar regression-based approach to handling time-to-event phenotypes, via simulations and with a focus on small to moderate sample sizes.

This work was motivated by a search for genes that modify the Duchenne muscular dystrophy (DMD) phenotype. DMD is an X-linked recessive disorder affecting ≈ 1 in 5,000 live male births (5, 6). DMD involves progressive muscle tissue loss with replacement by fat and fibrotic tissue, and is currently without a cure. Patients typically become reliant on wheelchairs by early to mid-adolescence, but some maintain ambulation substantially longer, and age at loss of ambulation (LOA) is an important clinical indicator of disease progression. A great deal is known about the gene (*DMD*) that causes DMD, including the fact that modifier genes influence the rate of disease progression in a DMD mouse model (7, 8); evidence for modifiers exists in humans as well (9–12). The discovery of modifier genes in humans has implications both for therapeutics and for the design of DMD clinical trials. Thus far there have been two published GWASs for LOA, based on sample sizes of 170 (11) and 253 (12) individuals, respectively.

Using data from the United Dystrophinopathy Project, a multisite consortium (13–15), we are engaged in a search for modifier genes under a GWAS design. The sample currently comprises ≈ 400 DMD patients, with ongoing efforts to increase the sample size to ≈ 800 DMD patients. (These patients are conservatively selected to exclude mutations in *DMD* itself that are known or suspected to affect disease course.) There is one binary covariate in the model: steroid use prior to LOA, which is known to increase LOA by about 3 years on average. The immediate motivation for the current work is to inform design decisions regarding the analysis of our DMD data, while illustrating issues arising for GWAS in smaller data sets which could be of relevance to other studies as well. The paper can also serve to illustrate key features of the PPLD for GWAS investigators unfamiliar with the method.

The remainder of the paper is organized as follows. In 2 (Methods) we present (2.1) the simulation methods and (2.2) data analytic methods used in what follows. In 3 (Results) we begin (3.1) by considering the distributions of the TE-PPLD and the CPH p-value under the null hypothesis H_0_ of “no SNP-trait association,” under various conditions that can affect those distributions. We then (3.2) consider the behavior of both statistics under a variety of models under the alternative hypothesis H_A_ of “SNP-trait association.” In (3.3) we briefly contrast the PPLD’s use of Bayesian sequential updating to confirm findings with the standard requirement of independent replication, again with a particular focus on small sample sizes.

Finally, in 4 (Conclusions) we perform a final experiment in which we loosely mimic an entire genome scan in an initial data set, following up at any findings in a separate data set. This section illustrates the practical implications of many of the results of the preceding sections, and can be read first, or even independently, for an overview of the effects the choice between CPH and the TE-PPLD can have on GWAS results.

## 2. Methods

In this section we describe (2.1) the simulation methods and models used to evaluate the behavior of the different statistical analyses, and (2.2) the statistical methods used to analyze the simulated data.

### 2.1 Simulation Methods

In order to mimic features of our DMD data set, our base model uses a sample size of N = 400 individuals, half of whom come from each of two covariate levels (*y* = 1, 2); for some purposes we also consider sample sizes of N = 200 (similar to some potential “replication” data sets currently available for DMD) and N = 800 (our target sample size over the next few years), as noted in context. We simulated only unrelated individuals. Details of the simulation methods follow. All simulations were conducted in Matlab using built-in cdf and inverse cdf functions.

Individuals were randomly assigned a genotype for a 2-allele locus (with alleles 1, 2) as a function of the Minor Allele Frequency (MAF). Unless otherwise noted, we set MAF = 0.5; for some comparisons we also vary MAF as noted in context. We generated genotypes under Hardy Weinberg Equilibrium (HWE). (The impact on the PPLD of violations of HWE and the impacts of of population stratification have been investigated previously (1) and are not further considered here.)

Under the null model H_0_ (no SNP-trait association; Model 1 in Table 1), event time *t_e_* was simulated via a random draw from a normal distribution *N_y=1_*(*μ, σ*) for individuals with *y* = 1, and *N_y=2_*(*μ* + 3, *σ*) for *y* = 2. Under the alternative model H_A_ (SNP-trait association), we simulated data under 7 different baseline Models 2-8 (Table 1), in which *t_e_* was randomly drawn from a mixture of normal (MoN) distributions in the form *N_y=1,k_*(*μ_k_, σ_k_*), for given genotype *k* = 11, 12, 22 and *y* = 1, and in the form *N_y=2,k_* (*μ_k_* + 3, *σ_k_*) for *y* = 2 (except for Model 8). Model 2 creates a simple additive mixture model for age-at-event while maintaining comparable x and s. *d.* at the population level. The remaining models vary effect size by increasing the genotypic variances (Model 3), introducing dominance (Models 4, 5), and by generating genotypic effects on variances as well as means (Model 6) or solely on variances (Model 7). Model 8 complicates the covariate effect. The models were chosen to illustrate a range of possible trait distributions, and are by no means intended to exhaustively cover what we might find in a real application. We simulated 1,000,000 replicates under Model 1, and 1,000 replicates per model under Models 2-8.

**Table 1.** Baseline simulation generating models. Generating parameter values were chosen to mimic LOA in the real DMD data set for uncensored individuals with no history of steroid use (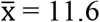, s. *d.* = 3.4). All normal distributions were left-truncated at 0 in order to preclude non-positive age at event. Models are shown on the standard normal scale for the *y* = 1 group. For Models 1-7, 3 years are added to the *y* = 1 means for the *y* = 2 group, as described in the text. For Model 8, *y* = 2 affects the means differently for the 3 genotypic groups in addition to affecting the variance; a comma separates the generating means (s.d.s) for *y* = 1, *y* = 2, respectively. These are the same generating models considered in (4).

| Model | $\mu_{11} (\sigma_{11})$ | $\mu_{12} (\sigma_{12})$ | $\mu_{22}(\sigma_{22})$ |
| --- | --- | --- | --- |
| 1 | 0 (1) | 0 (1) | 0 (1) |
| 2 | -0.5 (1) | 0 (1) | 0.5 (1) |
| 3 | -0.5 (1.25) | 0 (1.25) | 0.5 (1.25) |
| 4 | -0.5 (1.25) | 0.5 (1.25) | 0.5 (1.25) |
| 5 | -0.5 (1.25) | -0.5 (1.25) | 0.5 (1.25) |
| 6 | -0.5 (0.5) | 0 (1) | 0.5 (1.5) |
| 7 | 0 (0.5) | 0 (1) | 0 (1.5) |
| 8 | -0.5, 1.26 (1, 2) | 0, 1.76 (1, 2) | 0.5, 2.26 (1, 2) |

For each generating model, an individual was simulated based on a random draw of *t_e_*(*x_i_*) from the corresponding age-at-event (AE) distribution and an independent random draw of *t_o_*(*x_i_*) from an age-at-observation (AO) distribution. If *t_e_*(*x_i_*) < *t_o_*(*x_i_*), the individual was considered uncensored with failure time *t_fail_*(*x_i_*) = *t_e_*(*x_i_*); otherwise, the individual was considered censored with censoring time *t_cens_*(*x_i_*) = *t_o_*(*x_i_*). AO was simulated under a negative binomial distribution with *r* = 10, *p* = 0.4 in order to roughly mimic the censoring distribution in the real data. This yields a censoring rate ≈ 40%.

For some purposes, as noted in context below, we varied the baseline models. In considering robustness to the form of the underlying survival distribution, we also generated data (1,000,000 replicates per model under H_0_, and 1,000 replicates per model under H_A_) from Weibull (WB), Birnbaum-Saunders (BS), and Gamma (GM) distributions. This was done in each case by finding parameters of the distribution that matched the mean and standard deviation of the corresponding baseline model as shown in Table 1, and using these parameters as the generating values. We also considered covariate x genotype interactions; those models are described in context below.

### 2.2 Data Analysis Methods

In this section we give a brief overview of the TE-PPLD; for additional details see (4). We then describe the CPH analyses used in what follows, and we summarize some key differences between the TE-PPLD and CPH, which are relevant when comparing and contrasting results between the two methods. When referring to general features of Kelvin’s association statistic, we use “PPLD;” when discussing features that are (or may be) specific to the use of the PPLD with time-to-event data, we say “TE-PPLD.”

The PPLD is based on the Bayes ratio (BR), defined as

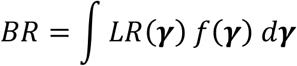

where LR is a likelihood ratio representing “trait-marker association” in the numerator and “no association” in the denominator (16), and the single integral stands in for multiple integration over the vector ***γ*** = *μ*_11_, *μ*_12_, *μ*_22_, *σ*_11_, *σ*_12_, *σ*_22_, the means and standard deviations of three normal distributions, one for each of the three SNP genotypes (17). For present purposes additional parameters of the likelihood are fixed as follows: recombination fraction *θ* = 0; standardized linkage disequilibrium (LD) parameter *D*’ = 1 (see (16)); admixture parameter *α* = 1 (see (18)); disease minor allele frequency (MAF) = SNP MAF. These simplifications allow us to model genotypic effects of the SNP itself (whether direct effects or indirect through LD) on either ***μ*** or ***σ*** or both. The underlying likelihood is based on the Elston-Stewart pedigree peeling algorithm (19) so that it can accommodate unrelated individuals as well as mixtures of pedigree structures. (This feature is helpful in our DMD study, because the dataset includes some pedigrees; however, we do not further consider it here.) The BR is proportional to a likelihood for the marker data conditioned on the trait data, and for reasons having to do with ascertainment corrections (20, 21) it is integrated as a unit, rather than separately in the numerator and denominator like a Bayes factor (22), using highly accurate non-stochastic numerical methods (23).

Let *π* be the probability that a randomly selected SNP is within detectable LD distance of a trait locus. The PPLD is a simple rescaling of the BR onto the (0,..,1) interval: 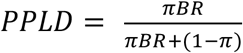. Thus PPLD < *π* indicates (some degree of) evidence in favor of H_0_, while PPLD > *π* indicates (some degree of) evidence in favor of H_A_; this remains true regardless of the value chosen for *π*. We set *π* = 0.0004, based in part on empirical calculations (1). By design, and in stark contrast with p-values, P[(PPLD > *π*) | H_0_] à 0 as N à ∞. In accumulating evidence for or against association across data sets, Bayesian sequential updating can be used by first multiplying the BRs across data sets and then applying the PPLD transformation to the resulting updated BR; we return to this in Section 3.2.4. All TE-PPLD calculations were done in Kelvin (20).

One limitation of Kelvin is that its models do not currently include any direct mechanisms for handling covariates. Our general approach to covariates is to preprocess the phenotype by performing regression analysis to make the covariate adjustments, and then to use the regression *residuals* as the input phenotype for subsequent analysis. In the context of linear regression, these residuals maintain the scale of the primary phenotype, and can be interpreted as estimates of how unusual is an individual’s phenotype given the individual’s covariate status. In the context of survival analysis, however, standard forms of residual (Martingale or deviance) do not maintain scale and do not have this “ordinary” interpretation (4). For this reason, we developed a new Ordinary Time-to-Event (OTE) residual, so-called because it maintains scaling vis a vis the primary phenotype and the interpretation of an ordinary linear regression residual, as a measure of how unusual the individual’s phenotype is given covariates. The OTE residuals then replace the underlying primary phenotype as input to TE-PPLD analysis.

In order to estimate OTE residuals for TE-PPLD analysis in what follows we use the procedures described and evaluated in (4). For each simulated data set, the estimated survival curve 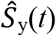, as a function of age *t*, is obtained via maximum likelihood estimation of a pair of 2-parameter Weibull distributions, one for each level of *y*, based on time-at-event for uncensored individuals and censoring time for censored individuals. OTE residuals for each individual are then calculated using the formula in (4).

PPLDs are reported to 2 decimal places for values ≥ 0.01 but to 4 decimal places for values < 0.01, in order to display whether very small values are greater than or less than the prior probability *π* = 0.0004, indicating evidence for or against association, respectively. In general we display results in terms of the TE-PPLD itself. However, because of the very low prior probability the PPLD scale is highly compressed at the low end. Thus for visualization purposes, particularly in considering the null distribution, we sometimes display log_10_BR instead.

CPH regression was performed using the built-in function of the Survival package in R (24). Regression analyses included the covariate *y* as well as genotypes as predictors; in considering models with covariate x genotype interactions (see section 3.2.3 below) we also include a *y* x genotype interaction term, as noted in context. Unless otherwise noted, we performed CPH analysis coding the genotypes to reflect the correct (generating) mode of inheritance (recessive, additive or dominant); under H_0_ we assumed an additive model. (By contrast, the PPLD does not require the user to specify a mode of inheritance.) CPH results are reported as P = −log_10_(p-value) for the genotypic coefficient unless otherwise noted, annotated as CPH-P.

Before proceeding to compare CPH with the PPLD, it is worth noting that the two approaches are in several respects incommensurate. The CPH p-value represents the probability of the data, or data more extreme, assuming H_0_, under the conditions imposed by the regression model; the TE-PPLD represents the posterior probability of H_A_, given the actual data only, under the assumptions described above. The p-value is not a measure of evidence strength (25), rather, it is considered significant when it crosses some preselected threshold. In GWAS contexts this threshold is conventionally set to 5×10^−8^, or P ≥ −log10(5×10^−8^) = 7.3, in order to adjust for multiple tests on a study-wide basis; in what follows we also consider a less stringent threshold of P ≥ 5.

By contrast, the BR is designed as a LR-based evidence measure (26–29). As a result, the PPLD provides an estimated rank-ordering of SNPs in terms of strength of evidence for or against trait-SNP association. Its calculation is not in itself a decision-making procedure, that is, there is no cutoff above which we declare significance; and, because it is not an error probability, it is not subject to multiple testing corrections. Nevertheless, with experience we have developed certain heuristics for prioritizing SNPs for further attention, with PPLDs ≥ 10% being of interest for follow up, and PPLDs ≥ 40% being of particular interest for follow up. (This rubric is context-dependent, however, much like the principle that it is fine to leave the house without a jacket whenever the temperature exceeds 70°F. This is a reasonable norm, but one which might be modified, if, say, one’s primary interest were in showing off a new jacket.)

## 3 Results

In what follows we evaluate the behavior of the TE-PPLD and CPH in application to GWAS analysis in the context of our intended genetic application, using simulated data. We have chosen the topics for the subsections to highlight some salient differences between the 2 methods in the GWAS setting, as well as to assist us in making practical decisions regarding how best to approach the analysis and interpret the results of our DMD study, or studies like the DMD study, with a focus on achievable sample sizes for relatively rare disorders. Except where specifically noted, we consider sample size N = 400.

### 3.1 Behavior of TE-PPLD and CPH regression under the null hypothesis

In this section we contrast the behavior of the TE-PPLD and CPH-P under the null hypothesis H_0_ of no association (Table 1, Model 1). Specifically, we consider: (3.1.1) fundamental differences in their sampling distributions; (3.1.2) effects of the form of the true underlying survival function S; and (3.1.3) the effects of varying MAF.

#### 3.1.1 Baseline behavior under H_0_

“*no association*” Figure 1 shows scatter plots of the the TE-PPLD compared with CPH-P as a function of sample size N, under H_0_. As can be seen, when there is no association the distribution of TE-PPLD moves leftward as the sample size increases, making large scores less and less likely. The distribution of the CPH-P is essentially constant as a function of sample size, as theoretically expected. In addition, the replicates with larger TE-PPLDs are not always the same as the replicates with larger CPH-Ps. As previously noted, the mathematical frameworks underlying calculation of CPH-P and the TE-PPLD are different, and this leads to different results not only in terms of the scales of the two statistics, but also in terms of rank-ordering.

**Figure 1.**
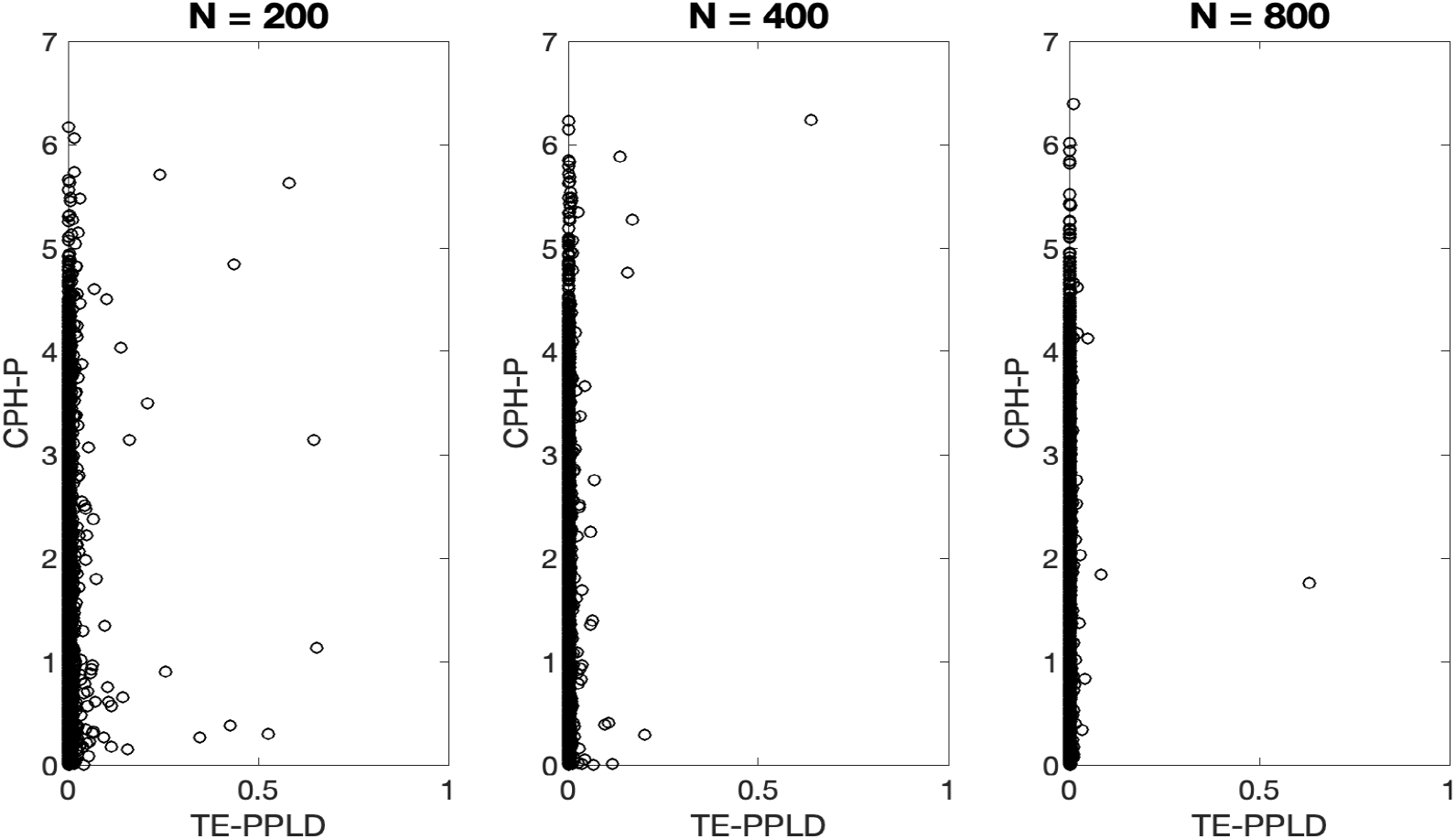
Comparative behavior of CPH-P, TE-PPLD under H_0_ as a function of sample size N. Shown here are scatter plots of the CPH-P and TE-PPLD distributions at three different sample sizes N, across 1,000,000 replicates generated independently for each N under H_0_: “no SNP-trait association.”

#### 3.1.2 Effects of changing the underlying time-to-event distributions under H_0_

In the previous section the generating distribution for time-to-event was normally distributed, as described above. Here we use 3 additional generating models: WB, BS and GM (see 2.1 above). Figure 2 compares the sampling distributions of each statistic across the different generating models. In this view, both log_10_BR (and therefore the TE-PPLD) and CPH-P appear to be relatively robust to the underlying form of S, although in both cases some pairs of generating distributions appear to differ at the upper end of the (respective) scales.

**Figure 2.**
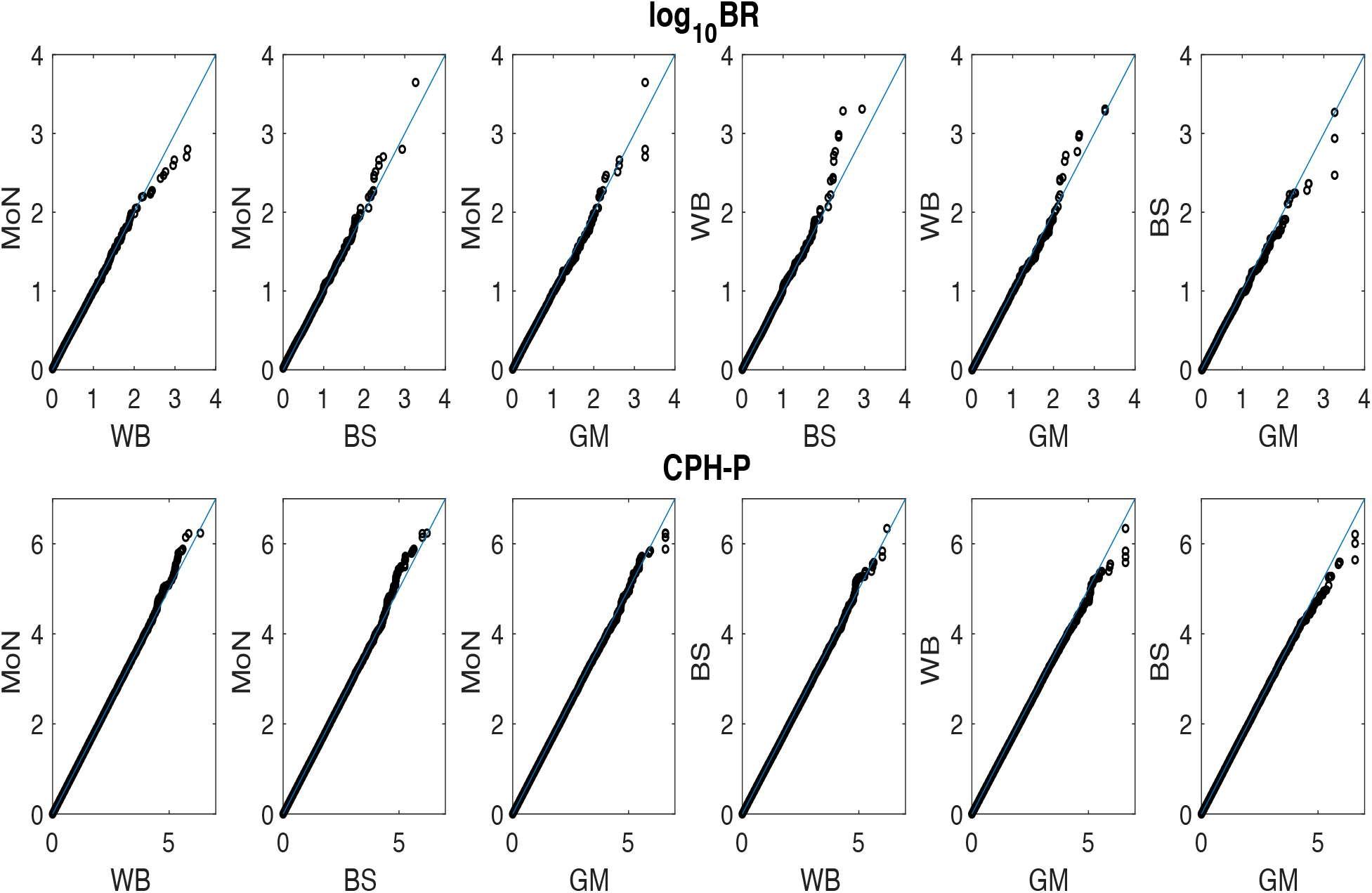
Comparative QQ plots for log_10_BR and CPH-P, respectively, as a function of generating form of S. Shown here are QQ plots for all possible pairs of generating distributions for S (Mixture of Normals (MoN), Weibull (WB), Birnbaum-Saunders (BS), Gamma (GM), for N=400, based on 1,000,000 replicates per generating condition.

In section 3.2.2 we will show some power calculations, and this would in principle require adjusting the significance thresholds under the different generating distributions were there an effect on the upper tail of the distribution. Adopting a significance threshold of P ≥ 5 returns 32 replicates above the threshold under the original MoN generating condition. (We use the lower significance threshold of 5 here because at this sample size there are no replicates with CPH-P ≥ 7.3 under any of these generating conditions; see also 3.2.4 below.) Table 2 shows the significance thresholds corresponding to the top 32 replicates under the other generating conditions, along with what would be corresponding cutoffs (i.e., demarcating the top 32 scores) for the TE-PPLD based on its null sampling distribution, were we to treat it as a test statistic in the conventional way. For both CPH-P and the TE-PPLD, the variation in thresholds across generating distributions is small, and with 1,000,000 replicates, very small differences cannot be estimated with high precision. Hence in what follows we utilize nominal Type 1 cutoffs without adjustment.

**Table 2.** Significance thresholds as a function of generating distribution. Estimated significance thresholds based on 1,000,000 replicates under H_0_, using CPH-P ≥ 5 under the Mixture of Normals (MoN) generating distribution, which demarcated the top 32 CPH-Ps, as a baseline. Thresholds corresponding to the top 32 SNPs are also shown for Weibull (WB), Birnbaum Saunders (BS) and Gamma (BM) generating distributions, as well as for the TE-PPLD.

|  | **MoN** | **WB** | **BS** | **GM** |
| --- | --- | --- | --- | --- |
| **CPH-P** | 5.0000 | 4.7567 | 4.7642 | 5.0293 |
| **TE-PPLD** | 0.0201 | 0.019 | 0.0202 | 0.0235 |

#### 3.1.3 Effects of lower MAFs

One issue of particular concern when using smaller sample sizes for GWAS is the effect of low MAF on the distribution of the test statistic under the null hypothesis. Regression analysis in general requires a sufficient number of individuals (say, at least 10-15) in the subsets created by division based on covariates: here that would entail requiring adequate numbers of individuals in each of the subgroups created by stratifying on genotype and the covariate *y* (3 x 2 = 6 subgroups). In small data sets many SNPs may fail to meet this bar; and the impact would be most pronounced under a recessive model. Here we consider the impact of lowering the MAF while assuming recessive inheritance for CPH analysis, and compare this with the corresponding impact on the TE-PPLD, for which the mode of inheritance is not specified (Figure 3). Note that with 400 individuals and a MAF = 10%, we only expect to see 4 individuals on average homozygous for the rare allele, which means only 2 on average in each covariate subgroup.

**Figure 3.**
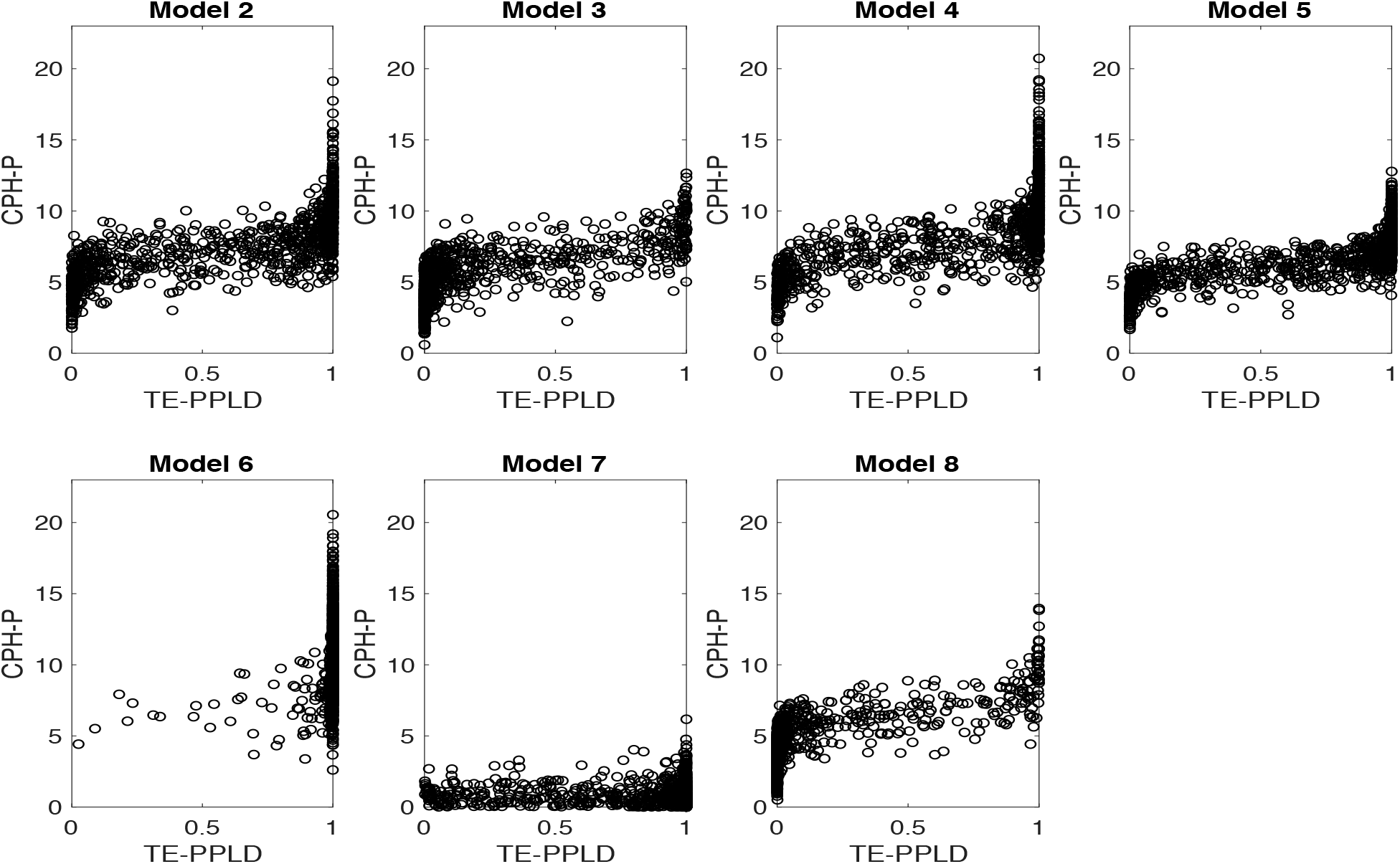
Effects of decreasing minor allele frequency under H_0_. Shown here are QQ plots comparing sampling distributions (N = 400) under H_0_ for minor allele frequency (MAF) = 0.4, 0.3, 0.2, 0.1 compared to the corresponding behavior with MAF = 0.5, for CPH-P and log_10_BR.

As the MAF decreases, the distribution of CPH-P becomes increasingly inflated, showing increasing (erroneous) larger P in favor of association. By contrast, for positive values (corresponding to TE-PPLD > *π*), there does not appear to be any inflationary effect on log_10_BR, at least until the MAF is quite small, and even then our interpretation of the results would not be materially affected: for instance, on the PPLD scale, the highest TE-PPLD is 0.96 for MAF = 0.1 and 0.64 for MAF = 0.5; in both cases, these would be clear cut “false-positive” results using our usual heuristics. The more notable effect on the TE-PPLD distribution is the progressive depression of log_10_BR for negative values as the MAF decreases, indicating increasingly larger (correct) evidence *against* association.

The built-in R routine for CPH returns NaN (“not a number”) under the recessive model when there are 0 individuals homozygous for the rare allele. Under the MAF = 0.1 condition, CPH returned a NaN for 18,242 SNPs. We confirmed that there is also little to no systematic bias against the null hypothesis when the TE-PPLD is applied to those replicates that were dropped by CPH (mean PPLD = 0.0003 < *π* (s.d. = 0.0019; max PPLD = 0.17)).

Of course, power to detect association will also be very low for SNPs with very low MAF, because there will be insufficient variability in genotypes to detect anything. It is common to drop SNPs with MAF below some threshold (say, 1-3%), with the threshold set higher for smaller data sets. Thus there are separate reasons for ignoring low MAF SNPs in the course of a genome scan. For CPH, one could additionally forego analysis under a recessive model in order to avoid this problem; even at MAF = 0.10, there were 951 SNPs with CPH-P ≥ 5 under the recessive model, but there were only 59 such SNPs under the additive model (and the additive results included the 18,242 SNPs dropped from the recessive analysis; note that this may also represent some inflation of scores under the additive model at low MAF, since we would expect to see on average 10 SNPs with CPH-P ≥ 5). But this of course risks missing a true recessive association, which has been suggested for some DMD modifier loci (30). The TE-PPLD, which does not incur the same upward bias under the null in small samples that plagues CPH under these conditions, does not force this choice.

### 3.2 Behavior of TE-PPLD and CPH regression under the alternative hypothesis

In this section we explore the behavior of TE-PPLD and CPH-P under generating models in which there *is* an association between SNP genotypes and the time-to-event phenotype. In 3.2.1 we show baseline comparisons for Models 2-8 (Table 1); in 3.2.2 we explore robustness to different forms of generating distributions; in 3.2.3 we consider an additional set of generating models involving epistasis, as described in that section. Finally, in 3.2.4 we consider challenges to independent replication as a gold standard for GWAS when only small to moderate sample sizes are attainable.

#### 3.2.1 Baseline results under the alternative hypothesis H_A_

“*SNP-trait association* Table 3 shows TE-PPLD and CPH-P results for the baseline alternative models (Models 2-8 in Table 1). Expected TE-PPLDs and CPH-Ps each vary as a function of generating conditions, with increasing means as sample size increases, as one would expect. Note that Model 7 involves effects on variances only; we would not expect CPH to detect association under this model. These baseline models were originally chosen in (4) to vary the mode of inheritance and the expected TE-PPLD while maintaining some reasonable ability to detect association at N = 400. In Table 3 these appear to be fairly strong effect sizes, in the sense that by N = 800 both methods are on average able to clearly detect association. However, recall that here we have used a generating MAF of 0.50; with lower MAFs average scores would be lower for both methods, in most cases appreciably so (see Section 4 below).

**Table 3.** Summary of sampling distributions of TE-PPLD, CPH-P, respectively, for the baseline models under H_A_. Shown here are the mean (standard deviation) of the sampling distribution, across 1,000 replicates per Model and sample size, of each statistic under each of the alternative models 2-8 from Table 1. For CPH *y* and genotype are included in the model as covariates; CPH analyses are run under the generating mode of inheritance (recessive, additive or dominant) per Table 1.

| Model | TE-PPLD |  |  | CPH-P |  |  |
| --- | --- | --- | --- | --- | --- | --- |
|  | N=200 | N=400 | N=800 | N=200 | N=400 | N=800 |
| 2 | 0.11 (0.23) | 0.60 (0.41) | 0.99 (0.05) | 4.28 (1.81) | 7.64 (2.48) | 14.33 (3.35) |
| 3 | 0.03 (0.12) | 0.22 (0.33) | 0.82 (0.32) | 2.99 (1.48) | 5.37 (1.99) | 9.95 (2.89) |
| 4 | 0.15 (0.28) | 0.68 (0.39) | 0.99 (0.05) | 4.63 (2.10) | 8.59 (2.97) | 16.08 (4.10) |
| 5 | 0.13 (0.27) | 0.64 (0.40) | 0.99 (0.07) | 3.61 (1.45) | 6.57 (1.88) | 12.37 (2.71) |
| 6 | 0.32 (0.38) | 0.98 (0.08) | 1.00 (0.00) | 5.37 (1.88) | 10.39 (2.69) | 20.00 (3.80) |
| 7 | 0.11 (0.22) | 0.83 (0.29) | 1.00 (0.00) | 0.72 (0.68) | 0.95 (0.80) | 1.51 (1.06) |
| 8 | 0.02 (0.09) | 0.14 (0.28) | 0.67 (0.39) | 2.67 (1.37) | 4.68 (1.92) | 8.51 (2.50) |

One salient feature of Table 3 is the large standard deviations across the 1,000 replicates per generating condition. Even though the generating model in each case represents a straight-forward genetic association model, and not, for example, a complex mixture of loci with different effects, nevertheless, both the TE-PPLD and CPH-P can vary widely from replicate to replicate in samples of this size; under mixture models standard deviations would be even larger. Figure 4 illustrates the extent of variability for each statistic on its own and in comparison with one another. We return to some implications of this level of variability in Section 3.2.4 below.

**Figure 4.**
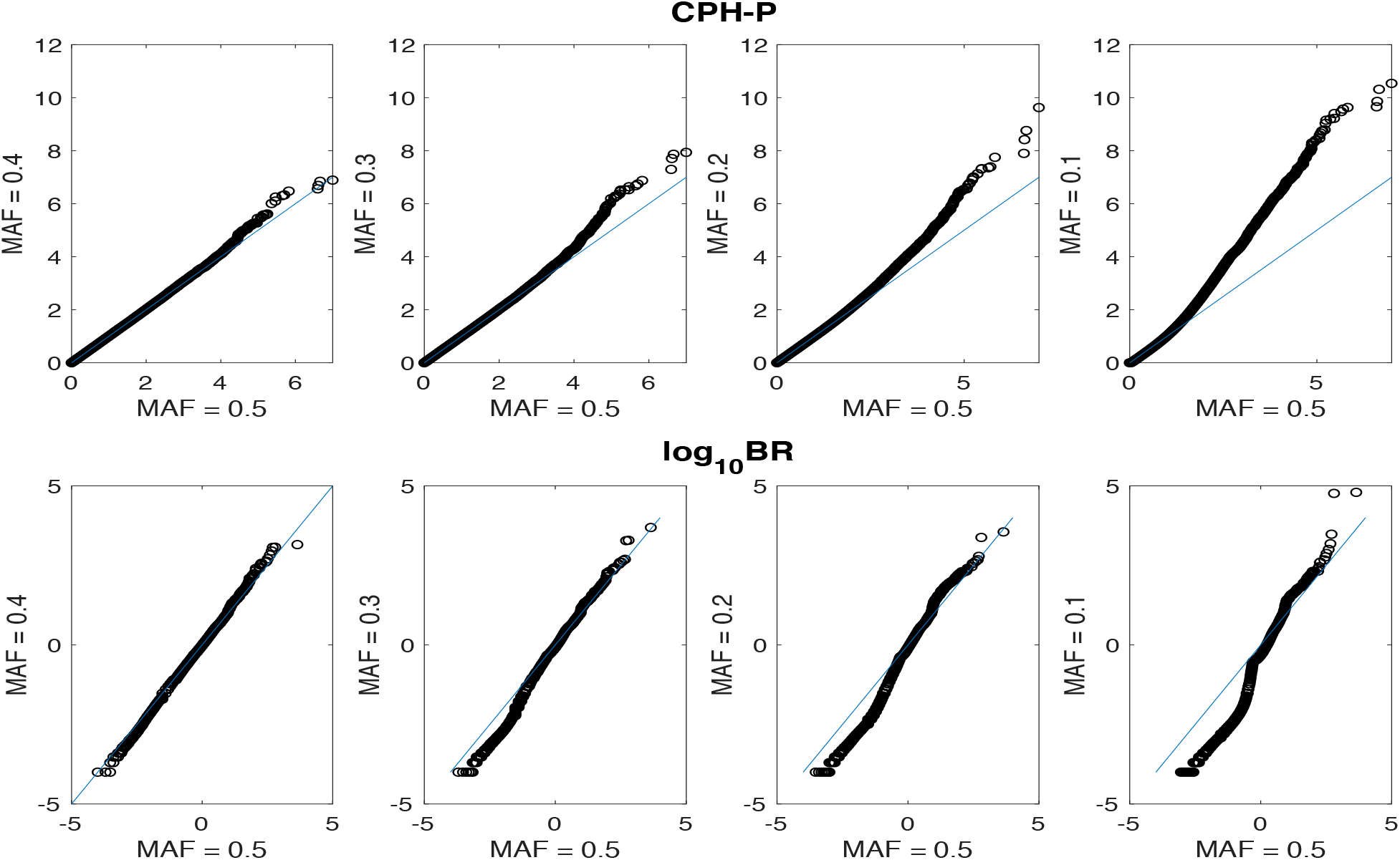
Comparative sampling variability of CPH-P and TE-PPLD under H_A_. Shown here are scatter plots for Models 2-8 from Table 1, with1,000 replicates per model (N = 400).

#### 3.2.2 Effects of changing the underlying time-to-event distributions under H_A_

Table 4 illustrates that under the alternative hypothesis the TE-PPLD is highly robust to the form of the underlying time-to-event distribution, across the range of distributions considered here. Thus neither the PPLD’s native “mixture of normals” assumption for a quantitative trait nor the use of the WB distribution for estimation of residuals complicates the interpretation of TE-PPLDs even when the underlying distribution violates these assumptions. By contrast, for some generating models the average CPH-P can drop considerably under some generating distributions.

**Table 4.** Robustness to true underlying time-to-event distribution under H_A_. Shown here are the mean (standard deviation) across 1,000 replicated generated under H_A_ (Models 2-8 from Table 1; N = 400). Results for the Mixture of Normals (MoN) distribution are repeated from Table 3 for comparison purposes; also shown are results for data generated under Weibull (WB), Birnbaum-Saunders (BS) and Gamma (GM) distributions. CPH regression included *y* and genotypes as covariates, and were run assuming the generating mode of inheritance (recessive, additive, dominant) for each model, as displayed in Table 1.

| Model | MoN | WB | BS | GM |
| --- | --- | --- | --- | --- |
| **TE-PPLD** |  |  |  |  |
| **2** | 0.60 (0.41) | 0.60 (0.41) | 0.56 (0.41) | 0.64 (0.40) |
| **3** | 0.22 (0.33) | 0.24 (0.34) | 0.21 (0.33) | 0.18 (0.30) |
| **4** | 0.68 (0.39) | 0.69 (0.39) | 0.71 (0.38) | 0.59 (0.42) |
| **5** | 0.64 (0.40) | 0.63 (0.40) | 0.59 (0.40) | 0.47 (0.41) |
| **6** | 0.98 (0.08) | 0.98 (0.10) | 0.96 (0.15) | 0.93 (0.19) |
| **7** | 0.83 (0.29) | 0.81 (0.31) | 1.00 (0.03) | 0.92 (0.20) |
| **8** | 0.14 (0.28) | 0.23 (0.34) | 0.61 (0.40) | 0.25 (0.35) |
| **CPH-P** |  |  |  |  |
| **2** | 7.64 (2.48) | 7.91 (2.55) | 5.34 (2.11) | 6.74 (2.35) |
| **3** | 5.37 (1.99) | 5.50 (2.05) | 3.78 (1.72) | 4.26 (1.84) |
| **4** | 8.59 (2.97) | 8.54 (2.84) | 6.04 (2.70) | 6.63 (2.78) |
| **5** | 6.57 (1.88) | 6.56 (1.86) | 4.59 (1.57) | 4.90 (1.59) |
| **6** | 10.39 (2.69) | 10.43 (2.62) | 6.28 (2.15) | 9.13 (2.61) |
| **7** | 0.95 (0.80) | 0.94 (0.83) | 0.48 (0.47) | 0.74 (0.67) |
| **8** | 4.68 (1.92) | 5.04 (1.88) | 5.87 (2.20) | 4.38 (1.73) |

**Table 5.**
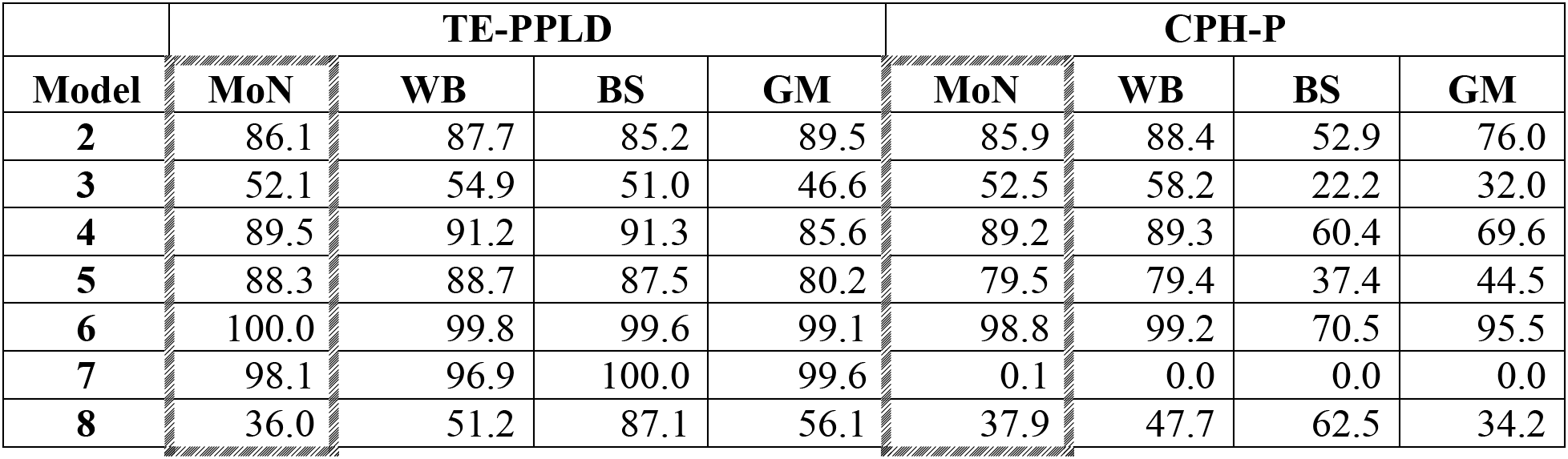
Comparative power of TE-PPLD and CPH-P analyses as a function of underlying survival distribution. Shown here is the estimated power to exceed a nominal CPH-P threshold of P ≥ 5, or the corresponding TE-PPLD threshold, when the underlying form of the generating model is Mixture of Normals (MoN), Weibull (WB), Birnbaum-Saunders (BS) or Gamma (GM).

| Model | TE-PPLD |  |  |  | CPH-P |  |  |  |
| --- | --- | --- | --- | --- | --- | --- | --- | --- |
|  | MoN | WB | BS | GM | MoN | WB | BS | GM |
| 2 | 86.1 | 87.7 | 85.2 | 89.5 | 85.9 | 88.4 | 52.9 | 76.0 |
| 3 | 52.1 | 54.9 | 51.0 | 46.6 | 52.5 | 58.2 | 22.2 | 32.0 |
| 4 | 89.5 | 91.2 | 91.3 | 85.6 | 89.2 | 89.3 | 60.4 | 69.6 |
| 5 | 88.3 | 88.7 | 87.5 | 80.2 | 79.5 | 79.4 | 37.4 | 44.5 |
| 6 | 100.0 | 99.8 | 99.6 | 99.1 | 98.8 | 99.2 | 70.5 | 95.5 |
| 7 | 98.1 | 96.9 | 100.0 | 99.6 | 0.1 | 0.0 | 0.0 | 0.0 |
| 8 | 36.0 | 51.2 | 87.1 | 56.1 | 37.9 | 47.7 | 62.5 | 34.2 |

Another way to look at this is from the point of view of conventional power calculations. Here we used nominal thresholds of P ≥ 5 for all CPH analysis, with corresponding thresholds for the TE-PPLD, as described in Section 3.1.2. As can be seen, the “power” of the TE-PPLD is affected very little by the underlying time-to-event distribution, across the range of generating distributions considered here. By contrast, CPH-P can suffer a loss of power, in many cases, a quite substantial loss, depending on the form of the underlying distribution.

#### 3.2.3 Genotype x covariate interactions

In studying the effects of modifier genes in the context of DMD, we are interested in the possibility that modifiers of the DMD phenotype might work by affecting response to treatment with steroids. The PPLD’s procedure for adjusting the residuals for covariate effects is done by “preprocessing” the data once, independently of genotype; by contrast, in a regression framework the covariate adjustment would be done separately for each SNP. In (4) we speculated that, as a result, the TE-PPLD might not be well powered to detect genotype x covariate interactions.

To investigate further, we simulated data under a variety of models involving covariate x genotype interactions as shown in Table 6. We note that the particular type of interaction we are considering here is a form of classical epistasis, in which the effect of genotype on the time-to-event may be masked by the absence of steroid exposure. Mathematically, this is only indirectly related to interaction in the usual statistical sense (31–33). It is, however, precisely the form of interaction of biological interest for the DMD study. In anticipation of section 3.2.4, where variable rates of covariate classification become important in the context of replication of findings, instead of simply setting N_*y*=1_ = N_*y*=2_ = 0.5, here we randomly draw the proportion *α* of individuals with *y* = 2 within each replicate from a *N*(0.7, 0.1) distribution.

**Table 6.** Generating models involving covariate x genotype interactions (epistasis) Models are shown on the standard normal scale. For each replicate, the proportion *α* of individuals with *y* = 2 is drawn from a *N*(0.7, 0.1) distribution; for all models, a value of *y* = 2 also adds 3 years on average to the mean time-of-event relative to *y* = 1, regardless of genotype.

| Model | $y = 1$ | | | $y = 2$ | | |
| --- | --- | --- | --- | --- | --- | --- |
| | $\mu_{11}(\sigma_{11})$ | $\mu_{12}(\sigma_{12})$ | $\mu_{22}(\sigma_{22})$ | $\mu_{11}(\sigma_{11})$ | $\mu_{12}(\sigma_{12})$ | $\mu_{22}(\sigma_{22})$ |
| **Epi 1** | 0 (1) | 0 (1) | 0 (1) | -0.5 (1) | 0.0 (1) | 0.5 (1) |
| **Epi 2** | 0 (1) | 0 (1) | 0 (1) | -0.5 (1) | 0.5 (1) | 0.5 (1) |
| **Epi 3** | 0 (1) | 0 (1) | 0 (1) | -0.5 (1) | -0.5 (1) | 0.5 (1) |
| **Epi 4** | 0 (1) | 0 (1) | 0 (1) | 0 (2) | 0 (1) | 0 (1) |
| **Epi 5** | -0.5 (1) | 0 (1) | 0 (1) | -0.75 (1) | 0 (1) | 0 (1) |
| **Epi 6** | -0.5 (1) | 0 (1) | 0 (1) | -0.5 (1.5) | 0 (1) | 0 (1) |
| **Epi 7** | -0.5 (1) | 0 (1) | 0 (1) | -0.75 (1.5) | 0 (1) | 0 (1) |

For the TE-PPLD, data were analyzed in two ways: (i) with each replicate (N = 400) “pooled,” that is, considered as a single data set; or (ii) dividing each replicate by covariate status and analyzing each of the two resulting data sets separately. For comparative purposes, we also analyzed the data under CPH, both without and with a covariate x genotype interaction term included in the model. Table 7 summarizes results. For the TE-PPLD, it is clear that in the presence of epistasis “pooled” analysis is much less effective than separate analysis in the *y* = 2 group, which is to be expected since the pooled group is a mixture of individuals, only some of whom represent any (detectable) genotypic effect. Also note that the TE-PPLD does an excellent job of distinguishing evidence for association from evidence against association, as reflected in the fact that for models Epi 1–4, in which there is no genotypic effect in the *y* = 1 group, TE-PPLD(*y* =2) > TE-PPLD(pooled); while for Epi 5-6, where there is some genotypic effect in the *y* = 1 group, TE-PPLD(*y* = 2) < TE-PPLD(pooled). By comparing the pooled results with the subset results we are therefore able to infer whether or not there is evidence of interaction. This works precisely because the TE-PPLD, by contrast with CPH-P, is able to indicate evidence for H_0_. Interestingly, under CPH analysis it seems preferable to not include the interaction term whether there is epistasis or not. Moreover, the interaction coefficient p-value is not a reliable indicator of whether or not epistasis exists.

**Table 7.** TE-PPLD and CPH-P results under epistasis models Epi 1 – Epi 7 from Table 6. Shown here are the mean (standard deviation) across 1,000 replicates, for the TE-PPLD and CPH regression. TE-PPLD results are computed in 2 ways : either with data “pooled” (N = 400) within each replicate, or with data subdivided based on *y* and analyzed separately in the 2 covariate groups. CPH analysis was run in 2 ways: without a *y* x genotype interaction term in the model (CPH-P Genotype = −log_10_(p-value) for the genotypic coefficient); or with an interaction term in the model. In the latter case we report both Genotype (g.t.), the value of P for the genotypic coefficient), and Genotype (interaction), the value of P for the interaction term. As above, CPH analyses were run assuming the generating mode of inheritance (recessive, additive, dominant) per Table 6.

| Model | TE-PPLD |  |  | CPH-P |  |  |
| --- | --- | --- | --- | --- | --- | --- |
| | *Pooled* | $y = 1$ | $y = 2$ | Genotype | Genotype (g.t.) | Genotype (interaction) |
| **Epi 1** | 0.08 (0.22) | 0.0001 (0.0002) | 0.21 (0.33) | 3.4 (1.9) | 0.5 (0.6) | 2.1 (1.4) |
| **Epi 2** | 0.30 (0.40) | 0.0001 (0.0002) | 0.62 (0.42) | 4.9 (2.8) | 0.6 (0.8) | 3.0 (1.7) |
| **Epi 3** | 0.22 (0.36) | 0.0002 (0.0022) | 0.57 (0.41) | 4.3 (1.9) | 0.6 (0.8) | 2.7 (1.6) |
| **Epi 4** | 0.02 (0.10) | 0.0001 (0.0006) | 0.04 (0.15) | 0.6 (0.6) | 0.5 (0.5) | 0.6 (0.6) |
| **Epi 5** | 0.33 (0.39) | 0.0046 (0.0407) | 0.25 (0.36) | 6.1 (2.4) | 2.0 (1.7) | 0.7 (0.7) |
| **Epi 6** | 0.10 (0.27) | 0.0020 (0.0112) | 0.05 (0.17) | 2.6 (1.7) | 1.9 (1.5) | 0.7 (0.7) |
| **Epi 7** | 0.35 (0.40) | 0.0039 (0.0359) | 0.25 (0.37) | 4.0 (2.2) | 2.0 (1.5) | 0.6 (0.6) |

Thus our earlier speculation in (4) that the TE-PPLD might not be useful for detecting covariate x genotype interactions appears to have been misplaced. For a binary covariate such interactions can apparently be detected by dividing the data set into two groups based on the covariate, then analyzing data with all of the data “pooled” (using *y*-adjusted residuals) and again separately in the two subsets; for the DMD application, where the interest is in a possible effect of steroid exposure (which would correspond to *y* = 2), this procedure would be applied only to the *y* = 2 group. SNPs with large TE-PPLDs in which TE-PPLD(pooled) > TE-PPLD(*y* =2) would then be indicative of association in the absence of an epistatic interaction, while SNPs with TE-PPLD(*y* =2) > TE-PPLD(pooled) would indicate an association involving interaction. In section 4 we revisit this approach in the context of a full genome scan.

#### 3.2.4 Independent replication vs. sequential updating based on small samples

As noted above, all of the generating conditions show strikingly high levels of variability across replicates in samples of the sizes considered here. This alone would suggest that clear-cut independent replication of a true signal might be unlikely, particularly when there might be only a few independent studies to use for comparison, each of which would also have a modest number of subjects, perhaps fewer than the initial study. To illustrate some of the issues involved in trying to replicate any DMD findings, we consider a situation we are likely to face, with access only to smaller replication samples for the time being.

In considering independent replication criteria based on p-values, there are many choices one could make regarding significance thresholds for the replication data set, and no clear answer as to which choice is correct. Here we use the NHGRI-EBI GWAS (https://www.ebi.ac.uk/gwas/) replication criteria: an association finding is considered to be replicated if either (Criterion 1) *both* of 2 studies gives P ≥ 5, or (Criterion 2) *pooling* the 2 data sets (or “mega-analysis”) gives P ≥ 5.

For purposes of illustration, we consider the additive Model Epi 1 (Table 6), and an initial data set of size N_*Init*_ = 400. Among the 1,000 replicates generated for Table 6 under this model, there were 190 with additive CPH-P ≥ 5. We selected 2 replicates to serve, respectively, as initial Candidate SNPs (CandSNP): one with CPH-P over the threshold of 5 (CandSNP#1_CPH-P_ = 5.70), and another with CPH-P meeting conventional genome-wide significance (CandSNP#2_CPH-P_ = 7.3). Note that the proportion of individuals with *y* = 2 was *α* = 65.8% in the CandSNP#1_CPH_ data set and 80.0% for CandSNP#2_CPH_. The higher value of *α* for the CandSNP#2 data set is an artifact of selecting the SNP based on a more stringent significance criterion, under conditions of variable *α*. We then attempted to replicate these signals in 1,000 independent replication data sets (RepSets) of size N = 200 under H_A_ (Model Epi 1), and in another 1,000 RepSets under H_0_ (Model 1, Table 1). Note that although the CandSNPs were drawn from a model involving a true association, here were are interested in our ability to *replicate* a finding: in this context it does not matter whether the initial finding is a true positive or a false positive; all that matters is the magnitude of the initial CPH-P, along with whether the RepSet itself comes from H_A_ or H_0_.

Just 3.7% of RepSets satisfied Criterion 1 under H_A_ (0% satisfied Criterion 1 under H_0_). That is, the probability of achieving statistical significance based on any given RepSet is negligible for this model at this sample size. Of course, with a larger RepSets this probability would increase; however, the large standard deviations, combined with the “winner’s curse” effect on *α* might still make clear-cut independent replication problematic.

Hence the only real possibility of satisfying the replication criterion under these circumstances comes from pooling the initial and replication data sets. Table 8 shows the percent of RepDS for which the pooled CPH-P exceeded replication Criterion 2. As can be seen, when the RepDS is generated under H_A_, our power to replicate the CandSNP is high under this Criterion for both CandSNPs #1 and #2. However, there is also a high false positive replication rate when following up with data generated under H_0_. Moreover, the larger the initial signal we are attempting to replicate, the more unreliable is Criterion 2, because the more the data in which the SNP was originally detected will dominate the pooled analysis, regardless of whether the replication data set itself supports association or fails to support association. Again, results would be different if the replication data set were larger than the initial one. Our point here is not to establish general power to replicate, but rather simply to illustrate some of the challenges of relying on replication to separate true from false positive findings in this setting.

**Table 8.**
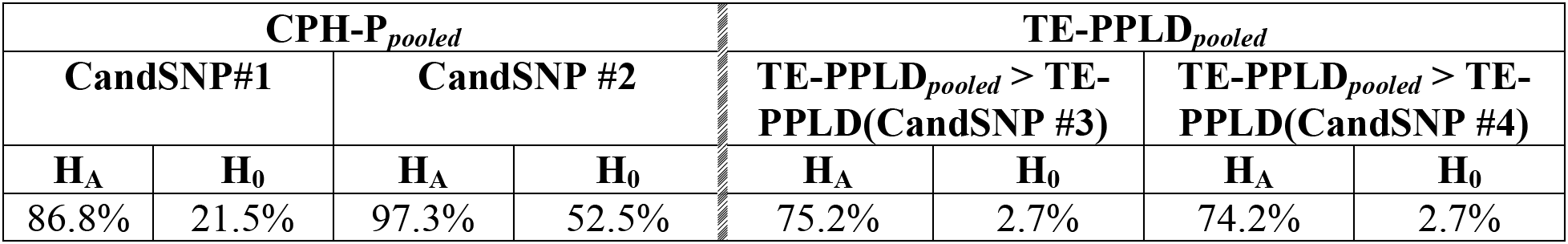
Probability of successful replication based on “pooling” initial and follow-up data sets, when following up on SNPs selected for moderate or high association signals. **CPH-P_*pooled*_**: Shown here are the proportion of pooled (N = 600) CPH-P exceeding the replication criterion of 5, when the initial data set (N = 400) is pooled with each of 1,000 replication data sets of size N = 200 each, generated under either H_A_ (Model Epi 1 from Table 6) or H_0_ (Model 1 from Table 1). CandSNP#1 selected based on CPH-P = 5.7 in the initial (N = 400) data set, CandSNP#2 selected based on CPH-P = 7.3. **TE-PPLD_*pooled*_**: Shown here are the proportion of pooled (N = 600) TE-PPLDs exceeding the initial TE-PPLD, when the initial data set is pooled with each of those same 1,000 replication data sets, generated under either H_A_ (Model Epi 1) or H_0_ (Model 1). CandSNP#3_TE-PPLD_ selected based on TE-PPLD = 0.53 in the initial data set, CandSNP#4_TE-PPLD_ selected based on TE-PPLD = 0.88.

| CPH-P<sub>pooled</sub> |  |  |  | TE-PPLD<sub>pooled</sub> |  |  |  |
| --- | --- | --- | --- | --- | --- | --- | --- |
| CandSNP#1 |  | CandSNP #2 |  | TE-PPLD<sub>pooled</sub> > TE-PPLD(CandSNP #3) |  | TE-PPLD<sub>pooled</sub> > TE-PPLD(CandSNP #4) |  |
| H<sub>A</sub> | H<sub>0</sub> | H<sub>A</sub> | H<sub>0</sub> | H<sub>A</sub> | H<sub>0</sub> | H<sub>A</sub> | H<sub>0</sub> |
| 86.8% | 21.5% | 97.3% | 52.5% | 75.2% | 2.7% | 74.2% | 2.7% |

For comparison, we selected separate replicates such that CandSNP#3_TE-PPLD_ = 0.53, which is large enough to satisfy our usual heuristics, and CandSNP#4_TE-PPLD_ = 0.88. Table 8 compares the pooled (N = 600) TE-PPLD with each of the initial CandSNP TE-PPLDs. What we see is that most of the time when the RepDS is generated under H_A_, the pooled TE-PPLD is larger than the initial CandSNP TE-PPLD, and the percentage of RepDSs with this feature is not dependent on the stringency of the criterion for selection of the CandSNP in the first place. Moreover, when the RepDS comes from H_0_, the pooled TE-PPLD is consistently << CandSNP TE-PPLD. Thus the TE-PPLD appears, at least based on this one set of generating conditions, to be a better approach to replication than CPH-P when replication involves simple pooling of the initial and replication data sets.

However, it could be argued that the pooled result is not really what we want, because it is driven to a large extent by the initial data set, which was selected specifically for the size of its signal at the CandSNP. This would be true for either CPH-P or the TE-PPLD, despite the very different implications in terms of sampling behavior under replication. When relying on p-values, this issue is hard to circumvent, because the only alternative is to use some version of Criterion 1, and as we have seen, even requiring less than conventional genome-wide significance can be very hard to achieve in a small follow-up data set. (One might consider some form of meta-analysis, but this also fails for very much the same reasons that pooling the data fails (34).)

In the PPLD framework, we do have an alternative, namely, to focus instead on the *accumulation of evidence strength* as new data are accrued, via the mathematically rigorous technique of Bayesian sequential updating (see Methods, above). The basic idea is quite simple: Because the PPLD can detect evidence both for H_A_ and also for H_0_ (which the p-value cannot), when a follow-up data set supports “association” then on average we will have BR > 1 (or equivalently, PPLD > *π*); whereas when the follow-up data set supports “no association” we will have BR < 1 (i.e., PPLD < *π*). Sequential updating ensures that when the replication data set supports H_A_ the PPLD increases upon consideration of the new data, while when the replication data set supports H_0_, the PPLD decreases.

Applying sequential updating in the current experiment, we find that 34.2% of RepDSs have TE-PPLD > *π* under H_A_, while 4% have TE-PPLD> *π* under H_0_. This tells us the probability (34.2%) that the sequentially updated TE-PPLD will correctly increase, relative to the initially selected CandSNP, regardless of the size of the initial TE-PPLD. While a 34% success rate may seem quite low, it is arguably a more realistic assessment of the likelihood of agreement between two datasets at any given associated SNP under the conditions simulated here.

## 4. Conclusions

In this paper we have evaluated the sampling behavior of the TE-PPLD in small to moderate samples sizes, and compared this behavior with the sampling behavior of CPH p-values, using simulations. We have noted a number of contrasts between the the TE-PPLD and CPH-P. Some of what we have found will be specific to time-to-event data, but most findings will apply to any application of the PPLD in the GWAS setting with small to moderate sample sizes.

We selected the sample sizes and topics covered here based on design questions facing a GWAS-based search for genetic modifiers of DMD, in order to inform our approach to analyzing and interpreting our own DMD data; and we have selected generating models to illustrate key points. We do not claim to have been exhaustive either in covering all possible topics or in covering all possible underlying genetic models. Nevertheless, the results presented above suggest several ways in which the TE-PPLD is a better choice than CPH in our setting.

In this final section we synthesize the implications of what we have found by loosely mimicking what might happen in a real GWAS for DMD modifiers. We assume 1,000,014 independent SNPs (no SNP-SNP linkage disequilibrium), 1,000,000 of which represent unassociated SNPs, simulated under H_0_; and 14 of which represent associated SNPs, simulated under H_A_ and generated 1 each from the 14 H_A_ generating models (Table 1, Models 2-8; Table 6 models Epi 1–Epi 7). We generated initial a single initial data set (InitDS) of size N = 400, and we followed up on selected SNPs in a single replication data set (RepSet) of N = 200. These simulations vary in 2 regards from those in the previous sections: (i) we generated *y* from a N(0.7, 0.1) distribution, as described in section 3.2.4, for all SNPs, separately in the InitDS and the RepSet; (ii) we used the MAF distribution from our actual DMD data set (Illumina Infinium Omni2.5Exome-8 v1.4, omitting SNPs with MAF < 3%; mean 0.23, s.d. 0.14). The RepSet data were generated from the same Model and using the same MAF that gave rise to each CandSNP in turn. Also here we consider a single replicate (at each sample size, N = 400, N = 200 respectively). This last experiment, therefore, is subject to “luck of the draw,” just as any single real study would be.

For CPH analysis we assumed additive inheritance for all SNPs. (We also repeated the experiment maximizing over the mode of inheritance at each SNP, but this approach resulted in far lower true positive rates; see footnote to Table 9.) We applied 2 significance criteria for selecting initial CandSNPs: either CPH-P ≥ 5.0, or CPH-P ≥ 7.3. We then followed up on all CandSNPs in the RepSet, again applying the NHGRI-EBI GWAS replication criteria as described above: a CandSNP was considered to be replicated if CPH-P(RepSet) ≥ 5 or if pooling the initial and replication data sets yielded CPH-P ≥ 5.

**Table 9.** True Positive Rates (TPR) for Initial Genome Scan (N = 400) and True Replication (Confirmation) Rates (N = 200), using various thresholds. TPR = True Positive Rate, or the proportion of all SNPs exceeding the threshold that represent true association; TRR = True Replication Rate, or the proportion of all SNPs with CPH-P meeting replication criteria that represent true association; TCR = True Confirmation Rate = the proportion of all SNPs with TE-PPLD > *π* in the replication data set that represent true association. CPH-P calculated under an additive model. (When maximizing over mode of inheritance (recessive, additive, dominant), at a threshold of 5 CPH-P picked up one additional true positive finding and 439 false positives, for TPR = 1%; when dropping SNPs with MAF < 10% the TPR improved only slightly, to 2%. At the higher threshold (7.3) the TPR was 4% and 10%, respectively, with and without the low MAF SNPs included.)

|  | Initial Genome Scan (N = 400) |  |  |  | Replication Data Set (N = 200) |  |  |
| --- | --- | --- | --- | --- | --- | --- | --- |
|  | Threshold | # of True Positive SNPs | # of False Positive SNPs | TPR | # of True Replications | # of False Replications | TRR |
| CPH-P | 5 | 2 | 29 | 0.06 | 2 | 4 | 0.33 |
|  | 7.3 | 1 | 1 | 0.50 | 1 | 0 | 1.0 |
|  | Threshold | # of True Positive SNPs | # of False Positive SNPs | TPR | # of True Confirmations | # of False Confirmations | TCR |
| TE-PPLD | 0.0430 | 4 | 29 | 0.12 | 3 | 0 | 1.0 |
|  | 0.10 | 3 | 9 | 0.25 | 3 | 0 | 1.0 |
|  | 0.40 | 3 | 2 | 0.60 | 3 | 0 | 1.0 |

For the TE-PPLD we considered 3 thresholds for determining CandSNPs: 0.0430, which corresponded in this data set to a CPH-P threshold of 5 under H_0_, and our usual heuristic thresholds of 0.10, 0.40. (Note that the threshold of 0.0430 is larger than the corresponding threshold of 0.0201 in Table 2. This is because Table 2 was generated with MAF = 0.5, while the current simulation involves variable MAFs, and it is consistent with the slight and largely inconsequential inflation of TE-PPLD scores under low MAFs as noted above.) We considered a CandSNP to be confirmed by the RepSet if the sequentially updated TE-PPLD > original (N = 400) CandSNP TE-PPLD, or in other words, if the CandSNP yielded TE-PPLD > *π* in the RepSet alone.

Table 9 summarizes the overall performance of the two methods. In the InitSet, using a threshold of 7.3 for CPH-P, 2 SNPs cross the threshold, one of which represents H_A_, and this was the CPH condition yielding the highest True Positive Rate (TPR) = 1/2 = 50%. The highest TPR for the TE-PPLD was 60%, which occurred when using the heuristic threshold of 0.4, and yielded 3 truly associated SNPs, compared to the 1 association correctly identified under CPH. It is also interesting to note that in the initial genome scan, using CPH-P criterion of 5 and the equivalent TE-PPLD, which yielded the same number of “false positive” signals by design, led to the correct identification of twice as many true positives under TE-PPLD analysis compared to CPH analysis. Note too that filtering out SNPs with MAF < 0.10 does not affect the number of true positive findings; however, it does reduce the number of false positive findings with the lower thresholds, from 29 to 23 for PPLD using 0.0430 (TRP = 14%), and from 29 to 17 for CPH-P using 5 (TPR = 11%).

CPH-P successfully replicated both CandSNPs crossing the threshold of 5 in the initial data set, including the 1 CandSNP that initially crossed the threshold of 7.3. However, replication was also seen for 4 of the 29 H_0_ SNPs initially crossing the threshold of 5, though not for the single H_0_ SNP initially crossing 7.3. Hence with the more stringent criterion for selecting CandSNPs, the True Replication Rate (TRR) was 100%; however, only 1 truly associated SNP was identified; at the lower threshold 2 truly associated SNPs were identified, but these made up just 2/6 SNPs satisfying the replication criteria.

By contrast, the TE-PPLD found confirmatory evidence for 3/4 H_A_ CandSNPs identified at the lower threshold of 0.0430 SNPs, and at none of the H_0_ CandSNPs, for a True Confirmation Rate (TCR) of 75%; while using the thresholds of 0.10 or 0.40 for selection of the initial CandSNPs, the TCRs were 100% in both cases (3/3 H_A_ SNPs confirmed and 0 H_0_ SNPs, at both thresholds). This means that one true positive SNP, identified in the initial data set, was not confirmed in the smaller, lower power follow-up data set. Hence failure to confirm, even in the context of these relatively simple generating models, does not necessarily mean that the initial finding was a false positive. But overall, the TE-PPLD identified more truly associated SNPs than did CPH-P, and with 0 false-positive confirmations using a threshold of 0.10 or higher for selection of CandSNPs in the initial data set.

Also informative is a comparison of rank-ordering of the TE-PPLDs or CPH-Ps (additive) (Table 10), of the 14 SNPs generated under H_A_ among all 1,000,014 SNPs. In this particular replicate, Model 5 yielded the top score for either method. The 2^nd^ largest TE-PPLD was obtained under Model 4 and ranked 3^rd^ (that is, the 2^nd^ highest TE-PPLD was from H_0_), while the 2^nd^ highest CPH-P was ranked 30^th^ (Model 2). The 3^rd^ highest TE-PPLD was from model Epi 3 and ranked 5^th^; while the 3^rd^ highest CPH-P was also from Epi 3 but ranked 62^nd^. Thus even under these conditions, in which it was apparently quite difficult to detect signals at most of the H_A_ models, the “true” positives tended to cluster closer to the top of the TE-PPLD rankings than the CPH-P rankings.

**Table 10.** Rank Order of “true positive” SNPs among all 1,000,014 SNP. CPH assumed additive inheritance. See Tables 2, 6 for generating models.

|  | TE-PPLD | CPH-P |
| --- | --- | --- |
| Model 2 | 29 | 30 |
| Model 3 | 94,032 | 6,381 |
| Model 4 | 3 | 6,547 |
| Model 5 | 1 | 1 |
| Model 6 | 2,796 | 634 |
| Model 7 | 16,653 | 223,516 |
| Model 8 | 276,148 | 19,234 |
| Epi 1 | 567,490 | 70,276 |
| Epi 2 | 599,782 | 602,124 |
| Epi 3 | 5 | 62 |
| Epi 4 | 89,758 | 720,810 |
| Epi 5 | 4,376 | 193,179 |
| Epi 6 | 533,873 | 611,318 |
| Epi 7 | 922,202 | 895,294 |

As a final experiment, we attempted to use stratification on *y* to detect covariate x genotype epistasis using the TE-PPLD, per section 3.2.3 above, particularly hoping to detect Models Epi 1-4 (Table 6). We considered two ways of doing this: (i) Based on the initial N = 400 data set, we rescanned the genome separately in the *y* = 2 group, accepting as evidence of epistasis any SNP at which the *y* = 2 TE-PPLD crossed a threshold (0.0430, 0.10, or 0.40) and also for which TE-PPLD(*y*=2) > TE-PPLD(N=400), that is, in which the subset-specific PPLD exceeded the “pooled” (across *y*) PPLD. Table 11 shows the results. In both cases, the singe true positive result occurred for Model Int 3. Rescanning the entire genome in y = 2 did not detect any additional true positive signals, compared to following up only on those SNPs already detected in the pooled analysis, which is somewhat surprising given the results in Table 7; however, it did produce additional false positive signals, no doubt due to the smaller sample size. (Recall that under H_0_, the PPLD is a less reliable indicator of evidence in favor of H_0_ the smaller the sample size.) The highest true confirmation rate (0.50) occurred when performing the separate y = 2 analysis only at SNPs crossing the 0.40 threshold in the initial pooled analysis. It is also notable that we did not erroneously infer epistasis for any of the non-epistatic H_A_ models.

**Table 11.** True Positive Rates (TPR) for Detection of Epistasis using the TE-PPLD in initial N = 400 data set, at various thresholds

| Rescan Genome in $y = 2$ | | | | Consider only SNPs crossing threshold in pooled ($y = 1, y = 2$) analysis | | |
| --- | --- | --- | --- | --- | --- | --- |
| Threshold | # of True Positive SNPs | # of False Positive SNPs | TPR | # of True Positive SNPs | # of False Positive SNPs | TPR |
| **0.0430** | **1** | **16** | **0.06** | **1** | **5** | **0.17** |
| **0.10** | **1** | **14** | **0.07** | **1** | **2** | **0.33** |
| **0.40** | **1** | **2** | **0.33** | **1** | **1** | **0.50** |

### In Summary

We have illustrated several advantages of the PPLD over regression analysis in the context of GWAS with small to moderate sample sizes, both for identifying candidate SNPs and for confirming them in follow-up data sets (Table 12). We considered a range of models for genotypic effects on a time-to-event phenotype, and found that these tend to have low to moderate power in the sample sizes considered here, particularly using a realistic minor allele frequency distribution as would be found on a standard SNP array. In addition, these models showed very high variability across replicates, leaving a large role for chance both in terms of which truly associated SNPs can be detected in any given study and also in terms of our ability to find the same SNPs in follow-up data sets.

**Table 12.**
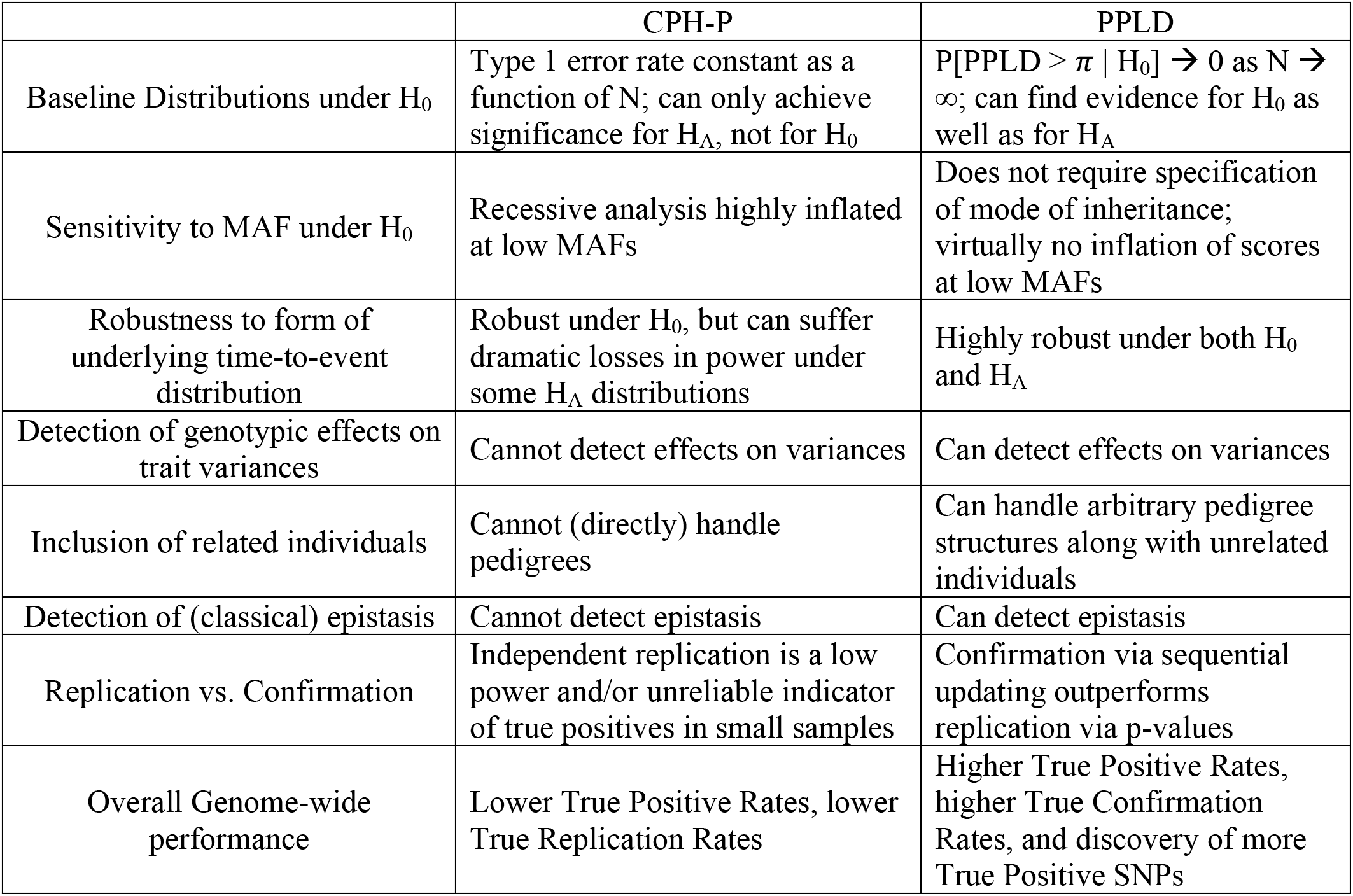
Summary comparisons between CPH-P and the PPLD

While there is no way to know in advance what sorts of models underlie true modifier effects in our DMD study, still, our results suggest overall that increasing sample sizes will be important, not only for the reliable detection of modifier genes but also for identification secondary effects such as covariate x genotype interactions. At the same time, however, our results support the overall reliability of TE-PPLD findings even in a data set of just N = 400; and they strongly support our contention that it is not necessary to aim for sample sizes in the the thousands or tens of thousands in order to reliably detect genes under the GWAS design, provided one applies statistical methods that are well-adapted to inference in small data sets.

## Data Availability Statement

This manuscript presents only simulated data. All computer code used in generating and analyzing the data is available at https://sourceforge.net/projects/kelvin-linkage-disequilibrium/files/.

